# Hitchhiking and epistasis give rise to cohort dynamics in adapting populations

**DOI:** 10.1101/106732

**Authors:** Sean W. Buskirk, Ryan Emily Peace, Gregory I. Lang

## Abstract

Beneficial mutations are the driving force of adaptive evolution. In asexual populations, the identification of beneficial alleles is confounded by the presence of genetically-linked hitchhiker mutations. Parallel evolution experiments enable the recognition of common targets of selection, yet these targets are inherently enriched for genes of large target size and mutations of large effect. A comprehensive study of individual mutations is necessary to create a realistic picture of the evolutionarily significant spectrum of beneficial mutations. Here we utilize a bulk-segregant approach to identify the beneficial mutations across 11 lineages of experimentally-evolved yeast populations. We report that most genome sequence evolution is non-adaptive: nearly 80% of detected mutations have no discernable effects on fitness and less than 1% are deleterious. We determine the distribution of driver and hitchhiker mutations in 31 mutational cohorts, groups of up to ten mutations that arise synchronously from low frequency and track tightly with one another. Surprisingly, we find that one-third of cohorts lack identifiable driver mutations. In addition, we identify intra-cohort synergistic epistasis between mutations in *hsl7* and *kel1*, which arose together in a low frequency lineage.

## INTRODUCTION

Adaptation is a fundamental biological process. The identification and characterization of the genetic mechanisms underlying adaptive evolution remains a central challenge in biology. To identify beneficial mutations, recent studies have characterized thousands of first-step mutations and systemic deletion and amplification mutations in the yeast genome (1, 2). These unbiased screens provide a wealth of information regarding the spectrum of beneficial mutations, their fitness effects, and the biological processes under selection. However, this information alone cannot predict which mutations will ultimately succeed in an evolutionary context as genetic interactions and population dynamics also impart substantial influence on the adaptive outcomes.

Early theoretical models assume that beneficial mutations are rare, such that once a beneficial mutation escapes drift, it will fix (3-5). For most microbial populations, however, multiple beneficial mutations will arise and spread simultaneously, leading to complex dynamics of clonal interference and genetic hitchhiking (6-9), and in many cases, multiple mutations track tightly with one another through time as mutational cohorts (10-13). The fate of each mutation is therefore dependent not only on its own fitness effect, but on the fitness effects of and interactions between all mutations in the population. Many beneficial mutations will be lost due to drift and clonal interference while many neutral (and even deleterious) mutations will fix by hitchhiking. The influence of clonal interference and genetic hitchhiking on the success of mutations makes it difficult to identify beneficial mutations from sequenced clones or population samples. Identifying beneficial mutations and quantifying their fitness effects ultimately requires assaying each individual mutation independent of co-evolved mutations.

Here we present a comprehensive, large-scale survey that quantifies the fitness effects of 116 mutations from eleven evolved lineages for which high-resolution knowledge of the dynamics of genome sequence evolution is known. We describe a novel strategy for constructing bulk-segregant pools that enables high-throughput quantification of fitness effects of individual evolved alleles. We find that large-effect mutations in common targets of selection drive adaptation while deleterious alleles rarely reach appreciable frequencies. We report that most genome sequence evolution is non-adaptive: nearly 80% of detected mutations have no discernable effect on fitness. Furthermore, one-third of cohorts lack identifiable driver mutations, and the dynamics of these “driverless” cohorts can be explained by genetic hitchhiking alone. Through the extensive characterization of evolved mutations, we begin to explore the mechanisms responsible for the observed cohort dynamics and we identify one cohort that suggests that rare mutations and epistatic interactions represent evolutionarily-relevant genetic mechanisms of adaptation.

## RESULTS

### Fitness of eleven representative lineages from nine evolved populations

Previously we evolved nearly 600 haploid *MAT***a** populations for 1,000 generations in a rich glucose medium. These populations were divided between two strains and two population sizes. “RM” and “BY” strains differ by a single SNP in *GPA1* that modifies the benefit of mutations in the mating pathway—a trait that we have studied extensively previously (8, 14). Different population sizes were maintained by controlling the dilution frequency and bottleneck size. We selected clones from both RM and BY populations for which we previously followed the frequencies of all mutations at high temporal resolution, selecting only those evolved at the smaller population size (Ne ~10^5^) to maintain uniformity across our experiments in this study. Since the number and identity of mutations in each population is known, we sampled across the range of biological pathways under selection (i.e. mating pathway, Ras pathway, cell wall assembly/biogenesis) and captured clones that exhibit a range of genome divergence, from 3 mutations in BYS1A08 to 16 mutations in BYS1E03. In two instances (RMS1G02 and RMS1H08), clones were isolated from two co-existing subpopulations to investigate how particular mutations and their fitness effects impact the ultimate fate of competing lineages. We identify selected clones by their population and the timepoint from which they were isolated (for example, clone BYS1A08-545). The evolutionary dynamics corresponding to each of the eleven selected clones are shown in their entirety in Figs. S2 and S3. We measured the fitness of each clone against a fluorescently-labeled ancestral strain of the appropriate background (BY or RM). Fitness values ranged from 2.7% in RMS1G02-545 to 8.6% in RMS1D12-910 (Fig. S1 and Table S1).

### Novel strategy for constructing bulk-segregant pools

We developed a novel bulk-segregant approach to rapidly and efficiently generate large pools of *MAT***a** segregants that contain random combinations of evolved mutations (Fig. S2). Briefly, we backcrossed evolved clones to a *MAT*α-version of the ancestor, gene converted the mating-type locus, and sporulated the resulting *MAT***a**/**a** diploid by complementing the *MAT*α*2* gene on a plasmid. Our method has three key advantages over the commonly used yeast “magic marker” approach, which selects on auxotrophic markers driven by mating-type specific promoters (15). First, our strategy ensures that the segregant pool is strictly MAT**a** through the removal of the *MAT*α gene during pool construction. Second, the approach produces segregants that are nearly isogenic to the evolved strain, thereby avoiding undesirable genetic interactions with the magic-marker machinery. Finally, the method can be applied to any strain background without requiring the incorporation of the magic-marker machinery and coordination of auxotrophic reporters.

### Fitness effect of 116 mutations across 11 evolved lineages

For each of our eleven evolved clones we generated a pool of ~10^5^ haploid *MAT***a** segregants. We propagated each pool for 90 generations and determined the fitness of each mutation by sequencing to a depth of ~100 reads per site every 20 generations (Fig. S2). The background-averaged fitness is calculated as the slope of the *ln*(mutant allele count/wild-type allele count) versus generations. In addition, we determined the segregation of fitness within each of the eleven crosses by isolating 192 segregants from each pool and quantifying fitness using a flow cytometry-based competitive fitness assay for a total of 2,112 fitness assays on individual segregants.

Two representative lineages are shown in Fig. 1, and all eleven lineages are shown in Fig. S1. We analyzed single clones from populations BYS2D06 and RMS1D12, each isolated from Generation 910. The evolutionary history of population BYS2D06 shows that eleven mutations swept through the population as three independent cohorts (Fig. 1) spaced several hundred generations apart. Bulk-segregant analysis revealed the presence of three beneficial mutations (*gas1*, 3.4%; *ira2*, 2.7%; and *ste5*, 3.3%) in the clone BYS2D06-910, each driving a single cohort (Fig. 1). The fitness distribution of the individual segregants shows distinct modes that fall within the range of fitness bounded by the ancestral and evolved parental strains. Population RMS1D12 exhibits more complicated dynamics (Fig. 1). The clone RMS1D12-910 contains fourteen mutations grouped into six cohorts (Fig. S3). Bulk-segregant analysis identifies three beneficial mutations (*ira1*, 3.3%, *rot2*, 5.6%, and *mid2*, 2.1%), each driving a different cohort. Again, the fitness values of the individual segregants fall between the fitness of the two parental strains, though the distribution appears more bimodal and less continuous than the distribution of fitness among the BYS2D06-910 segregants. These two major modes likely represent those segregants with and without the 5.6% fitness-effect *rot2* mutation.

**Fig. 1.**
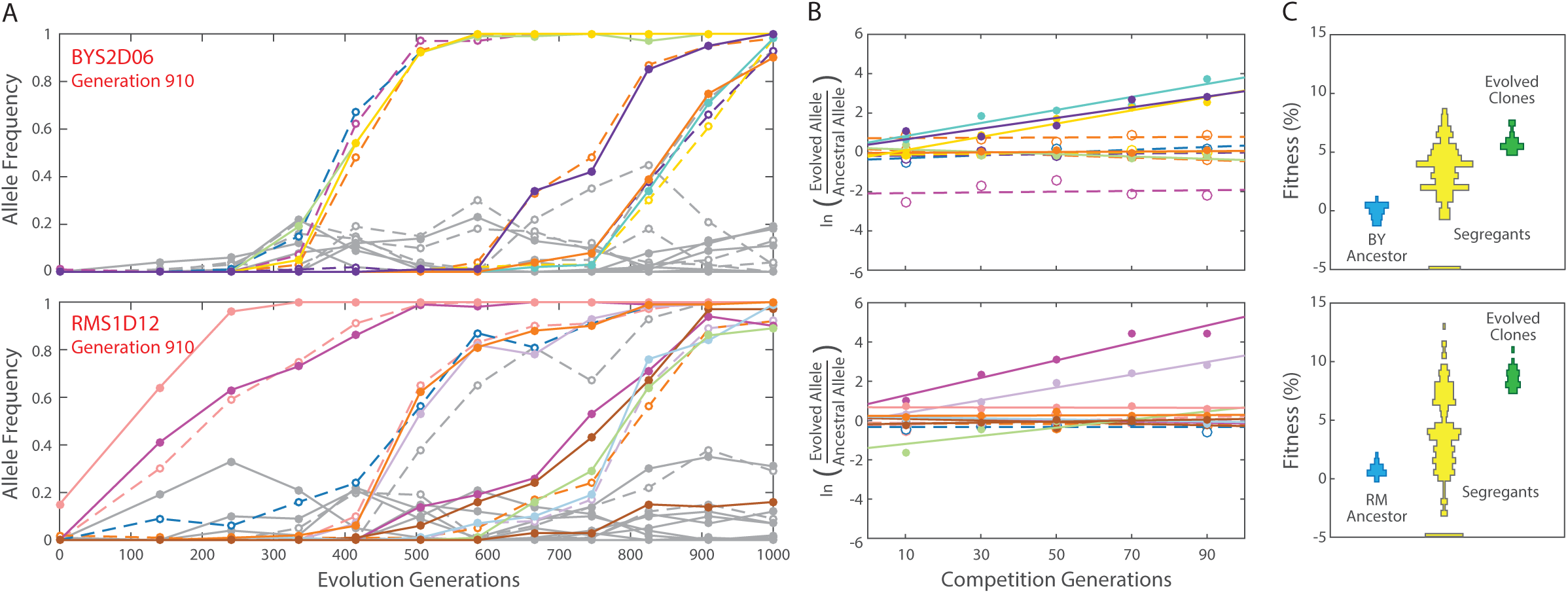
Genetic dissection of mutations from two evolved lineages. (*A*) Genome evolution of eachpopulation was previously tracked through time-course, whole-genome sequencing (10). An evolved clone was isolated from each population at defined time points. Each trajectory represents a unique mutation within the isolated clone (colored by chromosome), whereas gray trajectories indicate mutations detected in competing lineages within a population. (*B*) The background-averaged fitness effect of each evolved mutation is measured through a bulk-segregant fitness assay where a segregant pool is propagated in the selective environment and allele frequencies are tracked by whole-genome time-course sequencing. Fitness is calculated as the linear regression of the natural log ratio of evolved to ancestral allele frequency over time. The color scheme remains consistent between the evolutionary trajectories and bulk-segregant fitness assay. The dynamics of each mutation during the evolution experiment and the bulk-segregant fitness assay are in Fig. S1 and Table S2. (*C*) Individual clones isolated from a bulk-segregant pool are assayed for fitness against an ancestral reference in a flow cytometry-based competition to determine how fitness segregates in the cross (yellow). The fitnessdistribution of segregants derived from an ancestral cross (top panel: BY, bottom panel: RM) provides a baseline fitness in the absence of beneficial alleles. The fitness distribution of the individual segregants is compared to the fitness of the evolved clone from which they arose (green). The fitness for all 192 segregants from each of the eleven lineages is available in Table S3

Across the eleven lineages we measured the fitness of 116 of the 120 expected mutations. For four of the mutations we were unable to quantify fitness because all of the segregants in the pool contain the same allele due to gene conversion events during strain construction (see Discussion below). Of the 116 mutations across these eleven lineages, we find that 24 of the 116 mutations are beneficial, ranging in fitness from ~1% to ~10% effect. This indicates that most genome evolution is non-adaptive: roughly 80% of detected mutations are hitchhikers. As expected, all evolved clones possess at least one beneficial mutation. In addition, evolved clones contain up to 13 hitchhiker mutations (Fig. 2 and Table S2). Three mutations are reported as deleterious, though only one, a read-through mutation in the stop codon of *gcn2* in population BYS1D08, has greater than a 1% effect on fitness. We find no evidence of synonymous or intergenic mutations increasing fitness. When parsed by mutation type, driver mutations are enriched for nonsense and frameshift mutations compared to hitchhiker mutations (Fig. 3A). In contrast, synonymous and intergenic mutations are exclusively hitchhikers. This is consistent with adaptation driven by loss-of-function mutations in coding sequences (16).

**Fig. 2.**
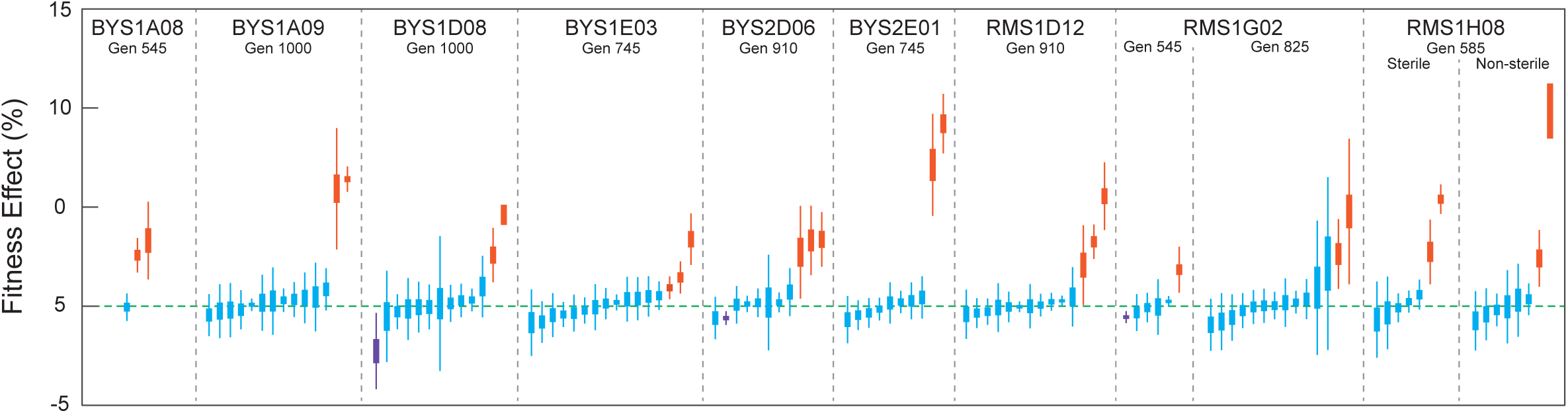
Comprehensive quantification of fitness effects. Background-averaged fitness effects for116 evolved mutations are quantified by bulk-segregant fitness assay. Mutations are separated by lineage of origin and ordered by mean fitness effect. Fitness effect is represented as the standard error of regression (thick lines) and 95% confidence interval (thin lines). Mutations are considered neutral (blue) if the confidence interval encompasses zero. Beneficial (orange) and deleterious (purple) mutations possess confidence intervals that fall completely above or below zero, respectively

### Mapping fitness effects to the dynamics of adaptation

Mutations often move through the populations as cohorts, synchronously escaping drift and tracking tightly with one another through time (10). Cohorts are a recent observation (10-13) and the evolutionary dynamics that drive their formation are yet to be explained. In general, mutations within a cohort are genetically and functionally unrelated, suggesting that cohorts are a random collection of mutations that accumulate while at low frequencies on a common background. A deeper understanding of these dynamics requires resolving the number of driver and hitchhiker mutations for a large sampling of cohorts. We used hierarchical clustering to objectively partition the 116 mutations into distinct cohorts based on correlated changes in allele frequency over time. Across the eleven selected lineages, we identify 31 distinct cohorts (Fig. S3). Each cohort contains up to ten mutations and each evolved lineage possesses up to six cohorts. Eighteen cohorts have a single driver mutation. Three cohorts each possess two unique driver mutations. These two-driver cohorts have twice as many hitchhiker mutations compared to single-driver cohorts (6.0 ± 2.0 and 3.1 ± 1.9, respectively, *P*=0.02, Wilcoxon Rank Sum, one-tailed), contributing to the overall positive correlation between number of beneficial mutations per cohort and cohort size (Fig. 3, *ρ*=0.70, Pearson correlation). Surprisingly, no beneficial mutations were detected within 10 of the 31 cohorts. These “driverless” cohorts typically contain few mutations; half are just a single mutation and only one is comprised of more than three mutations (Fig. 3B). We propose that driverless cohorts represent a form of genetic hitchhiking, where non-adaptive cohorts are “pulled” to intermediate frequency by the preceding cohort and/or “pushed” to higher frequencies during the sweep of the following cohort (Fig. S4).

**Fig. 3.**
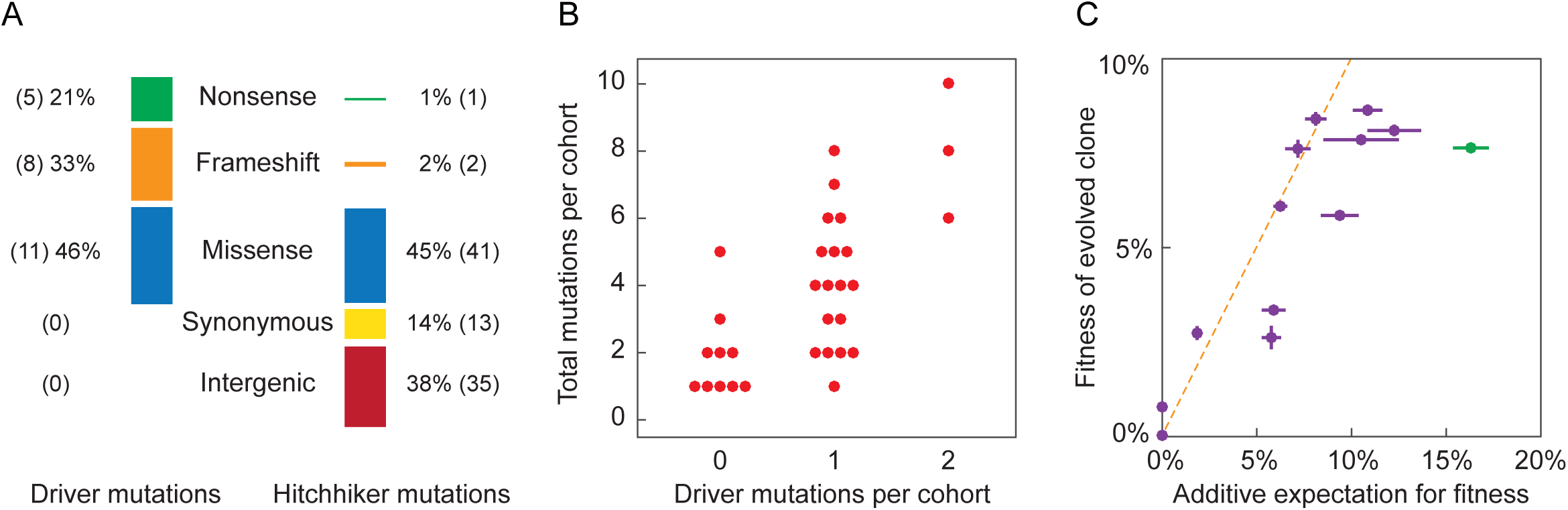
Mutational signatures, cohort composition, and additivity of fitness effects. A) Mutationswere divided into categories based upon their protein coding effect. The mutational signature of driver mutations is distinct from that of hitchhiker mutations (p<10^-9^; Chi-square). B) Hierarchical clustering identified 31 cohorts amongst the eleven evolved lineages. Cohorts vary in size from one to ten mutations and contain between zero and two drivers (Fig. S3). We observe a positive relationship between the number of drivers within a cohort correlates with cohort size (*ρ*=0.70; Pearson correlation). C)Fitness of all eleven evolved clones correlates with the sum of the fitness effects of their underlying evolved mutations, as quantified through bulk-segregant fitness assay. Vertical error bars reflect the standard error between replicate competitions of a common clone, and horizontal error bars reflect the propagation of error corresponding to the summation of individual background-averaged fitness effects. Deviation from the dashed line indicates non-additive genetic interactions. The BYS2E01-745 clone (green) deviates furthest from the expectation.

### Synergistic epistasis between *hsl7* and *kel1*, two genes whose protein products localize to sites of polarized growth

In the absence of epistasis, the fitness of an evolved clone is equivalent to the sum of the individual fitness effects of all constituent mutations. Indeed, we find a positive correlation (*ρ*=0.82, Pearson correlation) between the additive expectation of all mutations affecting fitness and the measured fitness of each evolved clone (Fig. 3C). Deviations from this additive expectation could indicate epistatic interactions between mutations that arose in the evolved lineage. One clone in particular, BYS2E01-745, deviates strongly from this additive expectation. Our bulk-segregant fitness assay identified two driver mutations in this lineage, *hsl7* (9.2%) and *kel1* (7.1%). The additive expectation (16.3%) is over twice the experimentally measured fitness of 7.6% (Fig. 3C). In addition, the BYS2E01-745 cross exhibits a highly asymmetric fitness distribution with a large proportion of low fitness segregants. (Fig. S1).

The dynamics of the BYS2E01-745 lineage are simple: an abrupt sweep of a single cohort of eleven mutations (Fig. 4A). However, the genetic basis of adaptation is unclear since BYS2E01-745 is the only lineage in this study without a mutation in a putative target of selection (10). Interestingly, the BYS2E01-745 lineage is enriched for mutations in genes whose products localize to the cellular bud and site of polarized growth (*P*=0.0012 and *P*=0.0016, respectively; GO Term Finder (17)). Further, all four genes (*hsl7*, *kel1*, *iqg1*, and *ccw12*), whose protein products localize to sites of polarized growth, contain either missense or frameshift mutations (Fig. 4A). Taken together, these data suggest that mutations in the BYS2E01-745 lineage may interact epistatically.

**Fig. 4.**
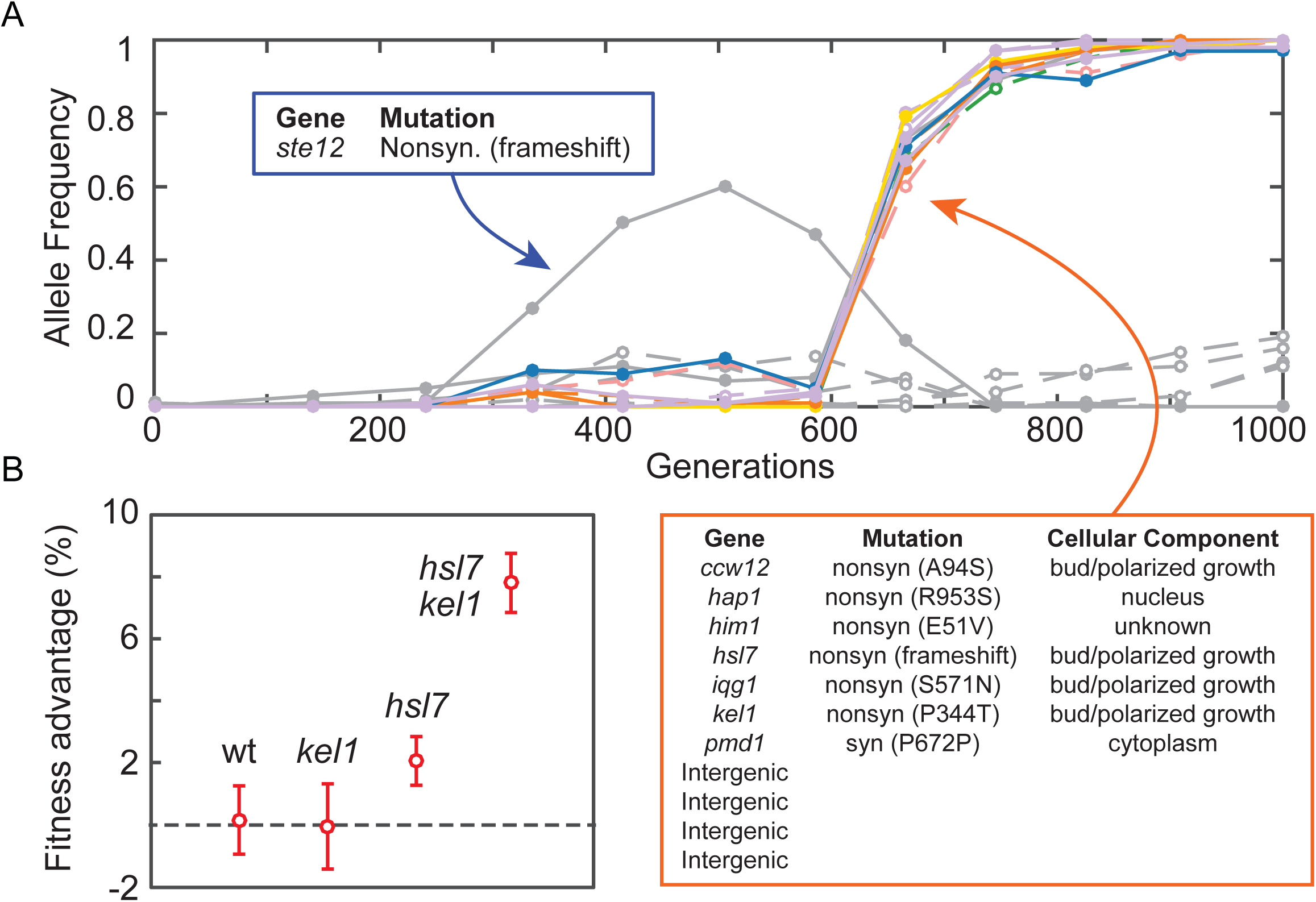
Adaptation mechanisms include rare mutations and epistatic interactions. A) Evolutionarydynamics of population BYS2E01 as tracked through whole-genome time-course sequencing (10). A beneficial *ste12* mutation (gray) outcompeted an 11-member cohort (colored) that is enriched for mutations in genes whose protein products localize to the cellular bud and site of polarized growth. B) Bulk-segregant individuals from the BYS2E01-745 cross were genotyped and assayed for fitness, producing a genotype-to-phenotype map. Two evolved alleles, *hsl7* and *kel1*, are associated with fitness gain. Shown are the average fitness and standard deviation of segregants when parsed by *HSL7*and *KEL1*alleles. The *kel1*mutation only confers a benefit in the *hsl7*background, and thefitness advantage of the *hsl7 kel1* double mutant is greater than the sum of the *hsl7* and *kel1* single mutants.

To test for epistatic interactions in the BYS2E01-745 lineage we genotyped all 192 segregants using low-coverage whole-genome sequencing (Table S4). Of the eleven mutations in the BYS2E01-745 lineage, only two loci significantly affect fitness: *HSL7* and *KEL1* (*P* < 0.0001; N-way ANOVA, Type III SS), corroborating the results of the bulk-segregant fitness assay (Fig. S1). Furthermore, the pairwise interaction between the *hsl7* and *kel1* alleles significantly impact fitness (*P* < 0.0001; N-Way ANOVA, Type III SS). The *hsl7* mutation confers a modest benefit in the *KEL1* wild-type background (~2.1%), while it confers a significantly larger advantage (~7.9%) when paired with the *kel1* allele (Fig. 4B). The *kel1* mutation is nearly neutral (-0.1%) in the *HSL7* wild-type background, but provides a substantial benefit in the *hsl7* background (~5.6%). As initially suspected from the bulk-segregant fitness assay (Fig. S1), the *IQG1* allele is absent from all individual segregants, the result of an endoduplication event that occurred during the construction of the parental diploid strain (Fig. S5). Through allelic replacement of *IQG1* in individual segregants, we found that the evolved *iqg1* allele has no effect on fitness. These results demonstrate that the rise of the BYS2E01-745 lineage is driven by intra-cohort epistasis between the *hsl7* and *kel1* mutations, which combine to produce a high fitness genotype, the fittest of the six BY evolved clones (Table S1). Detection and characterization of the *hsl7- kel1* interaction has revealed the only instance of synergistic epistasis to arise within a long-term evolution experiment outside of the well-studied Cit++ phenotype in *E. coli* (18-20).

## DISCUSSION

Recurrence-based models represent a widely-accepted statistical approach for inferring which genes and biological processes are under selection, by identifying targets that are mutated more often than expected by chance (21-25). Such probability-based methods, however, will inherently neglect rare driver mutations and are unsuitable for quantifying fitness as occurrence is not necessarily indicative of fitness effect. Many variables, e.g. mutational target size, can impact how often mutations in a particular target are detected. Indeed, we find no apparent relationship between the fitness effect that a driver confers and its prevalence (Fig. S6). For instance, both *ira1* and *ira2* provide similar effects on fitness (~2.7% each); however, *ira1* mutations were observed 23 times across replicate populations, yet *ira2* only once. Of our 24 driver mutations, our recurrence model did not detect six because they were mutated only twice; these include four modest-effect mutations (between 1 and 2%) as well as the two epistatic interactors, *hsl7* and *kel1*, in population BYS2E01 (Fig. S7).

The power behind identification of drivers through the bulk-segregant approach lies in the ability to screen lineages with numerous divergent loci for adaptive alleles. In only 1,000 generations, our populations fixed up to 19 mutations and in many cases, multiple cohorts of mutations existed simultaneously in the population leading to complex evolutionary dynamics. In addition, factors such as mutational target size, epistasis, genetic hitchhiking, and clonal-interference strongly influence the identity of mutations that arise over the course of evolution. Therefore, directly measuring the fitness effects of all mutations, as we do here, is necessary to unambiguously identify and quantify the fitness effects of driver mutations that could otherwise be missed by recurrence-based methods.

There is increasing evidence that large-effect beneficial mutations drive adaptation in microbial evolution experiments (1, 2) as well as in natural microbial populations (26) and clinical infections (27). Here we identify 24 driver mutations ranging from roughly 1% to 10% effect, consistent with populations evolving under the strong-selection strong-mutation paradigm in which small fitness effects (<1%) are unlikely to contribute to fitness. Because our populations are asexual it is possible for deleterious mutations to fix by hitchhiking on the background of strong driver mutations. We were surprised, however, to find that deleterious mutations are nearly absent from our dataset. Only three of 116 mutations were identified as deleterious, two of which are weak mutations (-0.6%) that could represent false positives. The lack of deleterious mutations is in contrast to an earlier study that found frequent hitchhiking of deleterious mutations in similar populations (12). We note that our use of 95% confidence intervals rather than standard error is more conservative, and may account for most of the discrepancy. We do have compelling evidence for at least one significant deleterious mutation: a stop codon readthrough of *gcn2* in population BYS1D08-1000. This mutation converts a stop codon (TAG) into a lysine residue (AAG), resulting in a predicted 26-amino acid extension of the C-terminus of the protein. The background-averaged fitness effect of this mutation is -2.26%, which is likely an underestimate of its cost since the mutant *gcn2* allele was not detected in generation 90 of the bulk-segregant fitness assay (out of the 107 reads that mapped to this position). It is unclear from our data if this *gcn2* mutation is deleterious in all backgrounds; therefore, it is possible that this mutation was beneficial on the background in which it arose.

Among the 116 mutations across the eleven evolved lineages in this study, we measured the fitness effect of multiple evolved alleles of the same gene, such as *ira1*, and multiple mutations in the same pathway, such as the mating pathway. The fitness effect of alternate adaptive alleles is remarkably consistent for the *ira1/2*, *gas1*, and *yur1* mutations, all of which differ by less than 1% in background-averaged fitness effect (Fig. S6). However, there are exceptions. A single neutral *ste12* mutation (frameshift at position 472) from the RMS1G02-825 lineage falls outside the narrow range (2.6%-3.3%) of the four adaptive mutations in the mating pathway. Additionally, the neutral *kel1-*Q107K mutation in the RMS1G02-825 lineage stands in contrast to the net-beneficial *kel1-*P344T mutation from BYS2E01-745 (Fig. S6), though we have shown that the latter is also neutral except in the presence of the co-evolved *hsl7* mutation. It is to be expected that established targets of selection acquire neutral mutations by chance so these evolved alleles may sweep by hitchhiking.

While common targets of selection may represent a substantial fraction of realized adaptive mutations, it is evident in this study and elsewhere (20) that some lineages may owe their success to the acquisition of a rare beneficial mutation or an assembly of epistatically-interacting mutations. Here, we identify synergistic epistasis between two mutations, *kel1* and *hsl7*, that arose within a single cohort. This observation leads to further questions surrounding the prevalence of such epistatic interactions, the context in which these interactions arise, and the extent to which epistasis, in addition to neutral genetic hitchhiking, gives rise to cohort dynamics.

In addition to quantifying fitness effects and identifying epistatic interactions, our data reveal a more detailed view of the dynamics of adaptation. Mapping driver and hitchhiker mutations to each of our 31 mutational cohorts shows that most cohorts consist of 1-2 driver mutations and 0-8 hitchhiker mutations. However, we also observe that approximately one-third of cohorts lack driver mutations. Visual inspection of the adaptive dynamics of these “driverless cohorts” reveals that their sweeps are intimately tied to other concurrent sweeps within the lineage. Driverless cohorts owe their success to hitchhiking, but unlike mutations within a cohort that arise synchronously from low frequency, mutations within driverless cohorts occur on the background of the preceding cohort during its climb. The ultimate fate of a driverless cohort is first tied to the sweep of the preceding cohort. If the preceding cohort is eventually driven to extinction by a competing lineage, the driverless cohort follows it into extinction (Fig. S4). Alternatively, if the preceding cohort fixes, the driverless cohort will remain at an intermediate frequency and its fate will be decided by the next selective sweep. We observe five instances of driverless cohorts fixing (Fig. S4). In each case this is a two-step process where a driverless cohort is first “pulled” by the preceding cohort and then “pushed” by the ensuing cohort. These “pull-push dynamics” explain a significant proportion, if not all, of the observed driverless cohorts. Based on our findings, we contend that driverless cohorts are prevalent across our evolution experiment and that the presence of a cohort does not necessitate the presence of a beneficial mutation. Our data therefore support three distinct modes of genetic hitchhiking: hitchhikers within a cohort, cohorts of hitchhikers “pulled” by a preceding driver mutation, and cohorts of hitchhikers “pushed” by a subsequent driver mutation.

Quantification of fitness can be complicated by intricate dependencies and interactions that have been shown to arise in laboratory evolution experiments due to nutrient crossfeeding or spatial structuring (28-32). One of our evolved clones (BYS1E03-745) exhibits negative frequency-dependent fitness when competed against an ancestral reference: the clone plateaued at frequency of 0.75 regardless of starting proportion (Table S1 and Fig. S8). We suspect the *erg11* mutation present in this clone since defects in ergosterol biosynthesis (or drug-based inhibition of the pathway) have been shown to result in negative frequency-dependence in yeast (28). We observed abnormal well-morphology *erg11*-containing cultures, with cells dispersed to the edges in addition to the bottom of the well, mirroring the morphology of the “adherent” A-type cells described previously. (28). Because our bulk-segregant pool contained random combinations of evolved mutations, both with and without the *erg11* mutation, we could successfully quantify the fitness effect of all mutations in this clone with the exception of the *erg11* allele itself, which maintained a frequency between 0.8 and 0.9 for the entirety of the 90-generation bulk-segregant fitness assay. Curiously, the *erg11* allele did not impart frequency dependence according to the adaptive dynamics of the evolving BYS1E03 population (10). Instead, *erg11* rapidly fixed in the evolution experiment. The *erg11* mutation may have been “pushed” through the population by the preceding *ira1*-driven cohort. In other words, the *ira1* allele may have masked the frequency dependence of the *erg11* allele.

This study demonstrates the power of experimental evolution to identify epistatic interactions. Much of our understanding of epistasis in budding yeast comes from the systematic analysis of double mutants (33). While informative for constructing large-scale genetic interaction networks, these studies have thus far been restricted to gene amplifications and deletions, which fail to capture a significant portion of the mutational spectrum. The observed *kel1*-*hsl7* interaction has not been identified by these systematic approaches, and several lines of evidence suggest that the interacting *hsl7* allele results from a rare mutational event. First, no other *hsl7* mutations were detected in any of the other 40 time-course sequenced populations (10) despite the BYS2E01-*hsl7* allele conferring a 2.1% advantage on the ancestral background. This *hsl7* allele encodes a truncated version of the protein that lacks only the C-terminal domain, which is phosphorylated by Hsl1p in cell cycle-dependent manner to relocate Hsl7p from the spindle-pole body to the bud neck (34). Deletion of *HSL7* is deleterious under a wide-range of conditions, including the rich glucose media used here (35, 36), thus our data suggests that the evolved *hsl7* allele bestows a novel function or alters an existing function. Extensive characterization of such rare beneficial mutations requires long-term, high replicate evolution experiments followed by comprehensive analysis linking genotype to phenotype. Likely due to their large target size, loss-of-function mutations dominate adaptive evolution experiments, though rare beneficial mutations and epistatic interactions may provide the raw material for molecular innovation in natural populations.

## METHODS

### Strain construction

We generated a *MAT*α strain with the following genotype: *ade2-1*, *ura3*∷*CAN1*, *his3-11*, *leu2-3,112*, *trp1-1*, *can1*, *bar1∷ADE2*, *hml*αΔ∷*LEU2*. This strain (yGIL737) differs from the ancestral strains from the original evolution experiment (8) in that it is *MAT*α, it has a loss-of-function mutation in *can1*, it has a wild-type copy of *CAN1* at the *ura3* locus, and it does not contain the NatMX cassette linked to *GPA1*, which is variable between the two ancestral strains (DBY15095/yGIL429 and DBY15092/yGIL432).

We crossed our *MAT***a** evolved clones to yGIL737 and, when necessary, we complemented sterile mutations in the evolved clones using the appropriate plasmid from the MoBY ORF plasmid collection (37). The resulting diploid strains were converted to *MAT***a**/**a** by a 3-hour galactose induction using a plasmid harboring a Gal-driven HO. *MAT***a**/**a** convertants were identified by the formation of shmoos following a 3h. αF-induction and were verified by PCR of the mating-type locus using primers *MAT*α-internal-F (5’ GCACGGAATA TGGGACTACT TCG 3’), *MAT***a**-internal-F (5’ ACTCCACTTC AAGTAAGAGT TTG 3’), and MAT-external-R (5’ AGTCACATC AAGATCGTTT ATGG 3’). Each *MAT***a**/**a** diploid was transformed with a *URA3* plasmid, pGIL071, which harbors the *MAT*α2 locus needed for sporulation. Note that using the full-length *MAT*α locus produces rare *MAT*α spores, presumably due to gene conversion at the *MAT* locus. Therefore the *MAT*α1 gene, which is required for *MAT*α mating but not sporulation, was not included on our plasmid.

### Generating segregant pools and isolating individual segregants

To create segregant pools, a single colony of each *MAT***a**/**a** diploid containing the plasmid-borne *MAT*α2 was inoculated into 10 ml of YPD and grown overnight to saturation. Overnight cultures were spun down and resuspended in 20 mL SPO++ (1.5% Potassium Acetate, 0.25% Yeast Extract, 0.25% Dextrose, supplemented with 1 × CSM-arg, Sunrise Science) and sporulated on a room temperature roller drum for 3-4 days until ~75% sporulation efficiency was reached. Sporulated cultures were spun down and resuspended in 200 µl H_2_O for a final volume of 500-750 µl. Ascus walls were digested by adding 5 µl of zymolyase (150 mg/mL) and incubating for 1h at 30°C. Next, 50 µl glass beads and 50 µl 10% Triton were added and the asci were disrupted by vortexing for 2 min. This was followed by an additional 40 min at 30°C and again by vortexing for 2 min. Disrupted asci were brought up to 5 ml with H_2_O and were sonicated using a microtip sonicator for four seconds at full power. A liquid hold was performed by inoculating 500 µl of the spore prep into 5 mL YPD and incubating for 24 hours at 30°C.

Segregants were isolated by plating onto 150 mm petri dishes containing solid BSP medium (CSM-arg, Yeast Nitrogen Base, 2% Dextrose, with 1 g/L 5FOA, 60 mg/L canavanine, and 100 mg/L ClonNat). To make segregant pools two milliliters of the spore preparation were plated onto duplicate BSP plates (resulting in ~10^5^ segregants across the two plates). After ~3 days of growth at 30°C, 5 ml of YPD was added to the first plate and a sterile glass spreader was used to remove yeast from the surface of the agar plate. This volume (2-3 ml) was transferred to the second plate and the process was repeated and the liquid was removed with a pipette and transferred to a 5 ml tube. To remove residual yeast cells, a second plate wash was done with 2.5 ml. This liquid from the second was (1-2 ml) was added to the same the same 5 ml tube for a total of ~4 ml. To isolate individual segregants, one milliliter of the spore prep was plated onto duplicate BSP plates. After ~3 days of growth at 30°C 192 colonies were picked and inoculated into 130 µl YPD in two 96-well plates. Plates were grown for 1−2 days at 30°C. All segregants were phenotyped for growth on 5FOA and canavanine, for the absence of growth on SC-ura, and for mating type. Segregants were stored at −80°C in 15% glycerol.

### Bulk-segregant fitness assay

For each bulk-segregant pool, seven replicate population were set up in a single 96-well plate. The bulk-segregant pools were propagated for 100 generations in conditions identical to the original evolution experiment (8). In brief, 132 µl cultures were maintained in YPD at 30°C and transferred to fresh media every 24 hours at a dilution of 1:1024 – identical conditions under which the populations were maintained during the evolution experiment. The assay lasted for a total of 10 days, corresponding to 100 generations. Following each transfer, the remaining culture was pooled across replicates (for a total volume of ~900 µL). Cells were pelleted and washed with 1 mL of H_2_O. Frozen pellets were stored at −20°C. Genomic DNA was prepared from the frozen pellets using a modified glass bead lysis method (10).

### Library preparation and whole-genome sequencing

Multiplexed genomic DNA libraries were prepared using a modified variation of the Nextera protocol (Baym *et al*. 2015) with the following modification: to conserve Nextera Index primers, the Index PCR was performed for 8 cycles, followed by amplification of the libraries via a 5-cycle Reconditioning PCR using Primers P1 (5’ AATGATACGG CGACCACCGA 3’) and P2 (5’ CAAGCAGAAG ACGGCATACGA 3’). Libraries were quantified by NanoDrop and pooled at equal concentration. The multiplexed pool was excised from an agarose gel (QIAquick^®^ Gel Extraction Kit, Qiagen) to size-select for fragments between 400-700 bp, and the collected fragments were analyzed by BioAnalyzer on High-Sensitivity DNA Chip (BioAnalyzer 2100, Agilent). The samples were run on an Illumina HiSeq 2500 sequencer with 250 bp single-end reads by the Sequencing Core Facility within the Lewis-Sigler Institute for Integrative Genomics at Princeton University.

### Sequencing data analysis

Raw sequencing data was split by index using a dual-index barcode splitter (barcode_splitter.py) from L. Parsons (Princeton University). To analyze the bulk-segregant fitness data we first generated a list of potential mutations that segregated within the bulk-segregant pool using our previous whole-genome, whole-population time-course sequencing data (10) to identify mutations within the evolved population at the time of clone isolation (Table S2). For each mutation, we generated four search terms corresponding to the ancestral and evolved alleles in both forward and reverse orientation. Each search term consisted of the known mutation as well as ten nucleotides immediately upstream and downstream of the mutation. In most instances, the search term was specific to a single locus within the genome. If the initial search term lacked absolute specificity in the reference genome, the search term was extended by five to ten nucleotides in each direction. For each FASTQ file, the number of reads containing the search terms was recorded, providing output in the form of ancestral and evolved allele counts at each locus. The search criteria required read accuracy/precision since only reads that possessed no mismatches/errors within the immediate vicinity of the mutation were counted. For comparison, the sequence reads were mapped to a corrected W303 genome (10) using BWA version 0.7.12 (38) with default parameters except “Disallow an indel within INT bp towards the ends” set to 0 and “Gap open penalty” set to 5. Mutations were then called using FreeBayes version v0.9.21-24-g840b412 (39) with default parameters for pooled samples. The two approaches were often harmonious (Fig. S9), and any significant discrepancies resulted due to FreeBayes calling the same mutation under several different call variants, particularly at the end of reads, an issue stemming from forced alignment to a designated ancestral reference.

The fitness effect of each mutation was calculated as the linear regression of the log ratio of allele frequency (N_evolved_/N_ancestral_) over the 100-generation experiment. Data corresponding to any time point at which either the ancestral or evolved allele was undetected were removed from analysis. Standard error of the regression and 95% confidence intervals were determined using MATLAB™ (MathWorks®). Mutations were classified as neutral if the confidence interval encompassed zero. Mutations with confidence intervals entirely above or below zero were characterized as beneficial or deleterious, respectively.

### Identification of additional allelic variants segregating in bulk-segregant pools

The genotyped bulk-segregant individuals from the BYS2E01 cross were analyzed for the presence of previously unrecognized genetic variants – alleles that differed between the *MAT***a** ancestor and *MAT*α parent, or low frequency mutations present in the evolved clone. VCF files corresponding to all BYS2E01 individuals were merged and scanned for calls that fit the following criteria: 1) the mutation is only called in a fraction of individuals, 2) the mutation, when called, is present near 100% in most clones, 3) the mutation is called at less than 100% in the individuals acknowledged to be non-clonal mixed samples, and 4) the mutation does not have another allelic variant. A total of seven genetic variants were identified and the sequencing data from the bulk-segregant pool fitness assay were analyzed using the search term approach (Fig. S10).

### *IQG1* allele replacement

All individuals in the BYS2E01-745 segregant pool contain the evolved allele of *iqg1*. To determine if the *iqq1* allele imparts a fitness effect, we replaced the *iqg1* mutation with the wild-type *IQG1* allele in 82 of the 192 segregant individuals. We synthesized a gBlock^®^ gene fragment (Integrated DNA Technologies) containing a guide RNA (AGAAAATATTATGAAGTTTT) targeting the *iqg1* mutation and the adjacent PAM sequence (CGG). The fragment was first cloned into the *URA3*-marked plasmid p426-SNR52p-gRNA-SUP4t (Addgene #43803). We then inserted the *HIS3* gene into the *BtgI* restriction sites in *URA3* creating a partial *ura3* deletion. The resulting *HIS3* gRNA plasmid, a *TRP1*-marked plasmid containing a constitutively expressed Cas9 (Addgene #43802), and a *URA3*-marked plasmid containing the wild-type *IQG1* allele (37) were co-transformed into 82 *iqg1* segregants. Transformants were selected on media utilizing the aforementioned auxotrophic markers, and all plasmids were eventually cured following verification of the allele swap. To screen for convertants, we utilized the *SspI* restriction site that was introduced through the evolved *iqg1* mutation (AGTATT → AATATT). Following PCR and *SspI* digest, transformants that yielded an intact 505 bp PCR product were presumed to be *IQG1* convertants. We performed Sanger sequencing on a subset of converted clones (6) to confirm the allele swap and ensure that no other mutations were introduced into the 505 bp region around the Cas9 cut site. Overall efficiency of allele replacement ranged from 15% to 85%.

### Flow cytometry-based competitive fitness assays

Fitness of the evolved clones and bulk-segregant individuals was determined in flow cytometry-based competitions as described previously (8, 10). Briefly, segregants were mixed 1:1 with a ymCitrine-labelled ancestral reference and passaged in YPD broth at 1:1024 dilution on a 24-hr cycle to mimic evolution conditions. Every 10 generations, an aliquot was transferred to PBS and assayed by flow cytometry (BD FACSCanto™ II). Cytometric data was analyzed by FlowJo™. Fitness was calculated as the linear regression of the log ratio of experimental-to-reference frequencies over the 40-generation assay. Extremely low-fitness clones, which presumably acquired a deleterious mutation, were not included in downstream analysis of the aggregate data. The clone isolated from population BYS1E03 displays negative frequency dependent selection due to a mutation in *erg11*. Mutations in the ergosterol pathway or chemical inhibition of Erg11p, give rise to spatial heterogeneity and negative frequency-dependent selection in our system (28).

## ACKNOWLEDGEMENTS

We thank Michael Desai, Andrew Murray, and members of the Lang Lab for their comments on the manuscript. This work was supported by The Charles E. Kaufman Foundation of The Pittsburgh Foundation.

## FIGURE LEGENDS

**Fig. S1: Genetic dissection of mutations from all eleven evolved clones.** (*A*) Genome evolution of each population was previously tracked through time-course, whole-genome sequencing (10). An evolved clone was isolated from each population at defined time points. Each trajectory represents a unique mutation, colored by chromosome, within the isolated clone, whereas gray trajectories indicate mutations detected in competing lineages within a population. (*B*) The background-averaged fitness effect of each evolved mutation is measured through a bulk-segregant fitness assay where a segregant pool is propagated in the selective environment and allele frequencies are tracked by whole-genome, time-course sequencing. Fitness is calculated as the linear regression of the natural log ratio of evolved to ancestral allele frequency over time. The color scheme remains consistent between the evolutionary trajectories and bulk-segregant fitness assay. (*C*) Individual clones isolated from a bulk-segregant pool are assayed for fitness against an ancestral reference in a flow cytometry-based competition to determine how fitness segregates in the cross (yellow). The fitness distribution of the individual segregants is compared to the fitness of the evolved clone from which they arose (green). Shown is the additive expectation of all driver mutations (blue) and all detected mutations (red) as measured by bulk-segregant fitness assay.

**Fig. S2: Construction of bulk-segregant pools and implementation of bulk-segregant fitness assays.** A) To uncover how evolved mutations impact fitness, an evolved clone is selected for bulk-segregant analysis. First, the evolved *MAT* **a** clone is crossed to a *MATα* version of the ancestor engineered to possess a functional *CAN1* gene and non-functional *can1* gene at unlinked loci. The resulting *MAT***a**/α diploid is heterozygous at all loci mutated in the evolved clone. The diploid is then mate-type switched to *MAT* **a**/**a** to ensure that only *MAT* **a** progeny are produced. To sporulate, the diploid is complemented with a plasmid harboring the *MATα2* gene, and isolation on canavanine-containing media selects against any unsporulated diploids. Each segregant contains a random combination of evolved mutations, and the pool, collectively, contains all possible genotypes. B) Each bulk-segregant pool is propagated in the selective environment for 100 generations, during which time allele frequencies change based on their fitness effect. Samples are collected every 20 generations and analyzed by whole-genome sequencing. C) The background-averaged fitness effect of a mutation is measured as the change in the natural log ratio of allele frequency over time. Beneficial mutations (red and blue) increase in frequency while neutral mutations (orange) remain steady over time.

**Fig. S3: Mutational cohort clustering and dynamics of selected evolved lineages.** Mutations were assigned into cohorts through grouping of evolved alleles based on a hierarchical clustering approach that leverages the known evolutionary dynamics. The heat maps reflect allele frequencies, and dendrograms display the distance and relationship between mutations. Mutations are colored by cohort designation, in accordance with both the dendrogram and evolutionary trajectories.

**Fig. S4: Driverless cohorts sweep due to their genetic association with neighboring cohorts.** Comprehensive analysis of evolved alleles revealed that a significant number of mutational cohorts do not possess any driver mutations. These driverless cohorts (red) often contain few mutations and exhibit evolutionary trajectories that mimic the preceding and ensuing driver-containing cohorts (gray). A driverless cohort can be pulled up by a previous cohort (e.g. Cohort B pulled by Cohort A in population RMS1G02, non-sterile clone), pushed up by an ensuing cohort (e.g. Cohort A pushed by Cohort B in population RMS1D12), or a combination of both (e.g. Cohort B pulled by Cohort A then pushed by Cohort C in population BYS1D08). The observed evolutionary dynamics of driverless cohorts can be explained without invoking adaptive mechanisms.

**Fig. S5: Sequence confirmation of the gene conversion event at the *IQG1* locus during construction of the BYS2E01 bulk-segregant pool.** To generate the BYS2E01 bulk-segregant pool, the evolved clone (*iqg1*) was crossed to the ancestor (*IQG1*). The resulting *MAT*a/α diploid exhibited the expected *IQG1*/*iqg1* genotype. However, after mate-type switching, the resultant *MAT*a/a diploid possessed only the *iqg1* allele, indicative of a gene conversion event. Following sporulation of the *MAT*a/a diploid, all haploid segregants possessed the *iqg1*allele.

**Fig. S6: Evolved mutations in common targets of selection confer similar fitness effects.** We compared the background-averaged fitness effects of mutations in common targets of selection, defined as genes or genetic pathways, which were mutated more than once within the eleven evolved lineages in this study. Thick bars refer to the standard error and thin bars refer to the 95% confidence interval of the background-averaged fitness effect of each mutation as determined through propagation and sequencing of each bulk-segregant pool (see Fig. 2). Both driver mutations (orange) and neutral mutations (blue) are represented. Each mutation is labeled by its predicted coding change (FS: frameshift, *: stop codon).

**Fig. S7: Fitness effect alone fails to predict the number of occurrences of each driver mutation.** The number of observed mutations in a given gene across forty replicate populations from Ref. (10) is not correlated to its background-average fitness effect (*ρ* = 0.11, *p* = 0.66, Spearman, two-tailed). For *gas1*, *kre6*, and *yur1*, the fitness values are the average of two measurements from independent populations. Similarly, for *ira1* the fitness value is the average of four measurements from independent populations. For *kel1* the fitness value is only from clone BYS2E01-745 and does not include the neutral *kel1* mutation in population clone RMS1G02-825. Open circles are driver mutations that were missed by our previous recurrence-based methods that identified putative driver mutations based on three or more observations across forty replicate populations (14). These include modest-effect mutations (~1-2%) as well as the large-effect interacting mutations, *kel1* and *hsl7*.

**Fig. S8: Evolved clone from population BYS1E03 exhibits negative frequency-dependent fitness.** Each evolved clone was assayed for fitness against an ancestral reference strain in seven replicate competitions. (*A*) In each competition, the evolved clone was seeded at four different starting frequencies (purple: 25%, green: 50%, orange: 75%, and red: 90%). The BYS2D06-910 competitions converge to a common ratio regardless of initial frequency, and subsequently its fitness (i.e. slope) decreases over time. In contrast, the fitness of BYS1E03-745 and all other evolved clones (Table S1) remains constant over time. (*B*) The instantaneous fitness values of clones BYS2D06-910 and BYS1E03 are calculated as the change in the log ratio of allele frequency across two consecutive timepoints (*n* and *n+10*). The fitness of clone BYS2D06 is inversely proportional to competition frequency (ρ=-0.72; Pearson correlation), whereas all other clones are independent of frequency, including BYS1E03 shown here (ρ=0.10, Pearson correlation).

**Fig. S9: Comparison of methods for determining allele counts within the bulk-segregant fitness assay.** Allele frequencies for the bulk-segregant fitness assay were determined through the analysis of Illumina sequencing data using two independent approaches: 1) BWA/FreeBayes and 2) our in-lab ‘search term’ approach. In both cases, the fitness effect of each evolved allele was calculated as the change in log ratio of allele frequency over time. Data from the first round of sequencing, which represents ~50% of total read coverage, was used to compare approaches. Manually investigation of the discrepancies validated the ‘search term’ approach, which was then used exclusively for all downstream analysis.

**Fig. S10: Newly discovered allelic variants have minimal impact on fitness.** Previously unknown allelic variants were identified through sequencing of the bulk-segregant individuals from the BYS2E01 cross. The frequencies of the seven discovered allelic variants were then tracked over time within all eleven bulk-segregant pools. Most variants are present in all pools, indicating a disparity between the *MAT*a ancestor and *MAT*α parent. An allele detected in only a fraction of the eleven pools (i.e. chrII_625980) was likely present at an intermediate frequency in the *MAT*a ancestral or *MAT*α parental culture. An allele specific to progeny from a single cross (i.e. chIV_593410) was likely a low-frequency evolved mutation. Most genetic variants appear neutral by the bulk-segregant fitness assay. Variants that depart from neutral are explained via linkage to a known driver mutation.

